# The presence of a marker associated with Pacific oyster resistance to OsHV-1 does not affect susceptibility to *Vibrio aestuarianus*

**DOI:** 10.1101/2024.11.25.625178

**Authors:** Liam B. Surry, Ben J. G. Sutherland, Spencer L. Lunda, Andrew H. Loudon, Konstantin Divilov, Chris J. Langdon, Timothy J. Green

## Abstract

The Pacific oyster, *Crassostrea* (*Magallana*) *gigas*, is an important species in aquaculture globally, but its production is threatened by pathogens including ostreid herpesvirus 1 (OsHV-1) and *Vibrio aestuarianus*. A genetic marker associated with resistance to OsHV-1 infection has been identified on chromosome 8 (Chr8) of the Pacific oyster genome. Marker-assisted selective breeding has used the Chr8 marker to produce oyster families with increased survivorship and resistance to OsHV-1 infection. The potential effect of the Chr8 marker on susceptibility to other pathogens remains largely unknown, but was recently associated with increased resistance of spat to *Vibrio coralliilyticus*. Here we assess the effect of the presence and allelic dosage of the Chr8 marker on the susceptibility of selectively bred juvenile Pacific oysters to *V. aestuarianus* infection. Sixteen families were produced with various Chr8 marker genotypes, and a *V. aestuarianus* disease challenge was conducted. Challenged oysters were individually genotyped at the Chr8 marker, and *Vibrio* susceptibility was evaluated among genotypes within and between families. Prior to the challenge trial, the Chr8 marker did not occur at the expected Mendelian ratios in families with heterozygous parents; in multiple families the homozygous alternate genotype occurred at a lower frequency than was expected. Mortality rates in the *Vibrio* challenge differed between families, ranging from 47.9% to 85.4%, but no association was observed between the Chr8 marker and survival to *V. aestuarianus* exposure. Therefore, we observed no pleiotropic effect (positive or negative) of the Chr8 marker on survivorship to *V. aestuarianus* at the evaluated life stage. Reduced-representation sequencing was used to genotype the challenged oysters and a genome-wide association study for *V. aestuarianus* survivorship was performed, but no significant associations were found, suggesting polygenic architecture for the trait.

## Introduction

The Pacific oyster *Crassostrea* (*Magallana*) *gigas* plays a prominent role in the aquaculture industry, accounting for more than 98% of all oyster production worldwide, with over five million metric tons produced annually (Azéma et al., 2017; Botta et al., 2020; Laurence et al., 2009; Okon et al., 2023). Native to the northwest Pacific Ocean, *C. gigas* has been introduced globally for aquaculture purposes because of its high growth rate and adaptability to various environments (Zhai et al., 2021). Aquaculture aims to supply aquatic-sourced proteins to meet human dietary requirements for the rapidly growing global population (Yang et al., 2022). However, the industry faces substantial challenges owing to pathogenic infections and outbreaks caused by viruses and bacteria (Yang et al., 2022). Since 2008, there has been a global increase in oyster mass mortality events, which has substantially decreased *C. gigas* production in Europe and Australia (Barbosa-Solomieu et al., 2015; Fuhrmann et al., 2019). For example, Pacific oyster production in New South Wales (Australia) decreased by 48% within three years of the first OsHV-1 outbreak. Decreases in production can cause significant increases in prices. In France, the price of oysters increased from € 4.93/kg in 2001 to € 7.78/kg in 2012, when the surviving spat from a 2008 mortality event reached market size (Fuhrmann et al., 2019).

Oysters do not possess adaptive immunity, and so resistance to pathogens cannot be enhanced through vaccination, but rather by methods such as immunostimulation, diet, and genetic selection (Divilov et al., 2021; Robinson & Green, 2020; Yang et al., 2022). Furthermore, oysters are typically cultivated in open environments within straits and bays, with complete contact with flow-through seawater, making it impossible to fully control diseases through the administration of disinfectants, antibiotics, or similar treatments. Quarantining is not feasible, and therefore, large-scale pathogen outbreaks can occur. Genetic improvement through selective breeding is considered to be the most practical approach to enhance disease tolerance and resistance (Green et al., 2023; Zhai et al., 2021).

Ostreid herpesvirus 1 (OsHV-1) is a pathogen that can cause large-scale mortality events in Pacific oysters. In 2008, a highly virulent form of OsHV-1 termed OsHV-1 microvariant (µVar), was detected in France (Segarra et al., 2010). Outbreaks of novel OsHV-1 μVars have subsequently occurred in Italy, the United Kingdom, Australia, New Zealand and the United States (de Lorgeril et al., 2018; Jenkins et al., 2013; Keeling et al., 2014; Burge et al., 2021; Divilov et al., 2023b). OsHV-1 affects oyster immunity and allows opportunistic bacteria, such as *Vibrio* species, to infect hosts (Siboni et al., 2024). OsHV-1 can infect all oyster life stages, especially larval and juvenile stages, resulting in up to 100% mortality (de Lorgeril et al., 2018). Therefore, substantial efforts have been made to limit the spread of OsHV-1. In 2018, a novel OsHV-1 µVar was detected in San Diego Bay, California, USA, posing a significant risk to oyster aquaculture on the west coast of North America (Burge et al., 2021; Thompson et al., 2024). Recent research using high-throughput sequencing indicated that this strain is different from the µVar detected in France as it has a 6 bp deletion in the microsatellite region instead of a 12 bp deletion, along with other differences (Pelletier, 2023). Given the threat posed by OsHV-1, substantial efforts have been made to implement strategies to mitigate the spread of the virus.

Extensive efforts in selective breeding programs have improved the OsHV-1 survivability of Pacific oysters in the USA and Australia as a potential mitigation strategy against OsHV-1 related mortality (Divilov et al., 2023; Kube et al., 2018). A quantitative trait locus (QTL) on chromosome 8 (Chr8) was identified as associated with an upregulated interferon regulatory factor (IRF) antiviral gene (i.e., *irf2*), and this is thought to have increased resistance to OsHV-1 infections (Divilov et al., 2021). Oysters with the alternate allele of this locus (hereafter referred to as the Chr8 marker) may be genetically primed to defend against viral infections. The Chr8 marker has been used in marker-assisted selection using a rhAmp genotyping assay (Beltz et al., 2018; Divilov et al., 2023), a fluorescence-based PCR method used to detect specific alleles at a locus. Selectively bred families carrying the Chr8 marker were produced by the Molluscan Broodstock Program (MBP) of Oregon State University. However, selective breeding may lead to fitness trade-offs (Divilov et al., 2024). For example, increased OsHV-1 resilience in offspring of oysters primed with an OsHV-1 viral mimic has been reported to co-occur with higher *Vibrio* infection levels in offspring (Robinson & Green, 2020). OsHV-1 resilience has also been correlated with decreased larval size and altered microbiome composition. Recently, families produced by the MBP via marker-assisted selection to produce either heterozygous or homozygous reference genotypes at the Chr8 marker were shown to have increased survivorship in heterozygous larvae and spat to *V. coralliilyticus* (Lunda et al., 2026). Collectively, this raises the question as to whether oysters bred for increased survival to OsHV-1 may have increased or decreased susceptibility to other bacterial pathogens, such as *Vibrio aestuarianus*.

Bacteria from the genus *Vibrio* are prevalent in marine environments, and several pathogenic strains threaten vertebrates and invertebrates (Wendling et al., 2014). *Vibrio* species can occupy a variety of niches, and include forms that are free-living, mutualistic, opportunistic, or pathogenic. Some *Vibrio* spp. can result in vibriosis, a disease that can cause mass oyster mortalities. Common species of *Vibrio* that are pathogenic to oysters include *V. splendidus*, *V. anguillarum*, *V. tubiashii* and *V. aestuarianus* (Wendling et al., 2014). In Europe, *V. aestuarianus* has caused recurrent mortalities since 2001, first in France and then in other European countries, including Italy, Ireland, and Spain (Mesnil et al., 2023). In recent years, the frequency of mortality events in France has increased from 30% in 2011 to 77% in 2013, when it has emerged as the primary pathogen identified during summer mortality events (Coyle et al., 2023). In British Columbia (BC), Canada, *V. aestuarianus* is one of the most common pathogenic *Vibrio* species and is emerging as a serious threat to oyster aquaculture of spat and adult *C. gigas* (Cowan et al., 2024). Considering the potential for the application of marker-assisted selection using the Chr8 marker on the west coast of North America and given that the effects of the Chr8 marker OsHV-1 survivorship allele on *V. aestuarianus* infection remain unknown, further investigation is required. This is particularly relevant considering the recent findings of an increase in resistance to oysters carrying the alternate allele of the Chr8 marker against *V. coralliilyticus* (Lunda et al., 2026). Additionally, although loci associated with increased survival to *V. alginolyticus* in Pacific oysters have been identified (Yang et al., 2022), the identification of loci associated with increased survival to *V. aestuarianus* would also be beneficial for selective breeding purposes.

In the present study, we evaluated the effect of allele presence and allele dosage of the Chr8 marker on the susceptibility to *V. aestuarianus* infection in a laboratory challenge. Given the functional differences in immunological responses to viruses and bacteria (Green et al., 2016), we hypothesized that there may be immunological trade-offs between resistance to viral and bacterial pathogens that may impact the resistance of oysters with the OsHV-1 survivorship allele to a bacterial infection. However, we may also see an increase in resistance, as was observed by Lunda et al. (2026) in infection trials with *V. coralliilyticus*. In either case, it will be important to characterize the response of oysters carrying the Chr8 marker alternate allele to *V. aestuarianus* infection. Furthermore, using the phenotypic data and samples from the laboratory trial, we performed double digest RAD-sequencing (Peterson et al., 2012) to conduct a genome-wide association study (GWAS) for *V. aestuarianus* survivorship.

## Methods

### Samples and crosses

Pacific oyster families with various genotypic states of the Chr8 marker (i.e. position 9,719,736 on Chr8; Divilov et al., 2023) were generated by Oregon State University’s Molluscan Broodstock Program (MBP; Oregon, USA). Spat from 16 families were provided to Vancouver Island University (VIU) to be used in experimental exposures to *V. aestuarianus*. Oysters (18 weeks old; > 4 mm long and < 6 mm wide) were transported to the Deep Bay Marine Field Station of VIU (Bowser, British Columbia (BC), Canada), held overnight in 12.5 °C flow-through seawater tanks, then transported to the Center for Shellfish Research (CSR) of VIU (Nanaimo, BC). Seawater and algae (*Chaetoceros muelleri*) samples were collected from Deep Bay Marine Field Station and transported to the VIU campus (Nanaimo).

Four of the 16 families had parents that were both heterozygous for the Chr8 marker (i.e., families F114-F117). The other 12 families had at least one parent with a homozygous reference genotype at the Chr8 marker locus (i.e., F101-F103, F105-F113; Table S1). These 12 families were the same families that were tested for resistance or susceptibility to *V. coralliilyticus* by Lunda et al. (2026). No families were produced by crosses with parents that were both homozygous for the alternate allele (i.e., the allele associated with increased survivorship to OsHV-1). Additionally, one family from the VIU oyster breeding program (Oyster Family Run 6, family 10; OFR6-10) was also included in the *Vibrio* challenge. OFR6-10 has not been selected for OsHV-1 resistance.

### *Disease challenge with* Vibrio aestuarianus

In western Canada, *V. aestuarianus* can cause farm mortality in juvenile and adult oysters during periods of elevated seawater temperatures (Cowan et al., 2024). Current laboratory methods for disease challenges in shellfish include injection, co-habitation, and bath treatments. Injecting oysters in the adductor muscle with a known amount of *V. aestuarianus* has been shown to provide reliable results (Green et al., 2016), but this method by-passes natural barriers of immunity (De Decker et al., 2011). Co-habitation potentially mimics a natural infection in the field (Dégremont et al., 2020, 2021) but this method can be unreliable in the timing and magnitude of mortality compared to injection or bath exposures (Nordmo et al., 1997). Therefore, here we conducted a bath exposure following procedures outlined by (Mackenzie et al., 2022) with a minor modification to include one oyster per well in the tissue culture plate to ensure that oysters are exposed to the same concentration of *V. aestuarianus*. It is worth noting here that factors known to affect disease risk to *V. aestuarianus* include oyster age, size, and exposure dose (Azema et al., 2015; Travers et al., 2017).

Forty-eight *C. gigas* from each of the 16 MBP families and the VIU family (total n = 816) were exposed to *V. aestuarianus*. An additional mixture of samples (n = 48) from multiple families was used as a non-exposed control group, treated in the same way as the exposed oysters but without exposure to *Vibrio*. For the *V. aestuarianus* exposure, *C. gigas* were placed in 12 well plates, where each well contained 3 mL seawater and one oyster, and then plates were acclimatized overnight in a temperature-controlled room at 16 °C.

A *Vibrio* suspension was prepared by inoculating *V. aestuarianus* (strain: BS032 2018) in two flasks containing TSB + 2% NaCl overnight in an orbital shaker (23 °C and 200 rpm). The cultures were then removed from the incubator, transferred into Falcon tubes, and centrifuged (5000 x g) at ambient temperature for 5 min. The supernatant was discarded and the cell pellets were resuspended in seawater. The optical density of the resuspended cells was measured at 600 nm using an BioSpectrometer and µCuvette® G1.0 (Eppendorf). A standard curve was prepared to estimate the amount of *V. aestuarianus* in the preparation. The concentration of the *V. aestuarianus* inoculum was also estimated by serial dilution and plating on tryptone soy agar with 2% NaCl, incubation at 23 °C for 48 h, followed by enumeration of colonies.

The exposure trial was initiated by adding 300 μL of the *V. aestuarianus* suspension to each well on each experimental plate (final concentration = 2.45 x 10^7^ CFU per ml). Sterile seawater was added to the control plate in place of the *V. aestuarianus* suspension. The temperature of the room was then increased to 23 °C for five days to simulate a marine heatwave event, and the temperature was reduced to 16 °C on the sixth day of exposure (Green et al., 2019). After *Vibrio* inoculation, 100 μL of algae (*C. muelleri*) was added to each well of all plates to encourage oyster feeding and uptake of *V. aestuarianus*. Morbidity was measured daily for six days, after which time the disease challenge ended. Oysters were considered moribund or deceased when their shell valves were gaping and failed to close after gently touching the oysters with a sterile toothpick. Oysters were placed into 25 mL Falcon tubes containing 100% ethanol, where whole individuals were pooled in storage tubes labeled with day of death and family. After six days, surviving oysters were preserved and stored in 100% ethanol. The ethanol-preserved oysters were stored at room temperature until DNA extraction was conducted. Mortality data were analyzed using IBM SPSS Statistics v28.0.1.0 (142) (IBM Corp., Armonk, NY) and R (R Core Team, 2026). Mortality distributions were assessed using a Kaplan-Meier survival analysis. A log-rank test was conducted to determine the differences in survival distributions among families, and a pairwise log-rank test was performed to determine which families had different survival distributions.

### Genomic DNA extraction

DNA extractions were performed on oysters from families F114-F117 (i.e., those with both parents heterozygous for the Chr8 marker) to extract total genomic DNA using a Monarch® Genomic DNA Purification Kit (#T3010L) according to manufacturer instructions for extractions from tissue. Briefly, oyster spat were removed from their shells using sterilized forceps, rinsed with deionized water to remove residual ethanol, then placed in a digestive solution containing lysis buffer and proteinase K (NEB). Samples were homogenized using a sterile pestle and digested at 56 °C for 2 h, after which an RNase incubation was performed. Genomic DNA was purified through an extraction column as per manufacturer’s guidelines and eluted in 35 μL molecular-grade water held at 60 °C. DNA quantification was performed via absorbance and Qubit fluorimetry using a BioSpectrometer and µCuvette® G1.0 (Eppendorf). Samples were checked for purity using A260/A280 and A260/A230 ratios on with the BioSpectrometer, and for quality using 1% agarose gel electrophoresis (see Supplemental Information). Extracted genomic DNA was then stored at -20 °C.

### Chr8 marker OsHV-1 resistance allele dosage

The rhAmp assay (Design ID: CD.GT.ZKJS3200.1; Integrated DNA Technologies) targeting a SNP with G/A (REF/ALT) alleles at position 9,719,736 bp on Chr8 (Divilov et al., 2023) was used to genotype the OsHV-1 resistance locus (i.e., Chr8 marker) in all individuals from the *V. aestuarianus* challenge trial in families F114-F117. The assay was conducted on a CFX96 Touch Real-Time PCR Detection System (BIO-RAD) using the manufacturer’s instructions for a 5 μL reaction volume in a 96-well plate (Table S2). Samples were normalized to 5 ng/μl in molecular-grade water and 10 ng (2 μL) of each sample added to each well, where each sample was run in duplicate. A negative control well was added to each plate, where instead of sample DNA it contained only molecular-grade water. The assay was previously confirmed to work on gBlocks as per manufacturer’s recommendations, and samples with known genotypes were used as positive controls in this study. The fluorophore dyes FAM and VIC were used for genotyping the reference (REF) and alternate (ALT) alleles, respectively, and to call genotypes as homozygous reference, heterozygous, or homozygous alternate.

Results of the rhAmp assay were exported from the instrument in text format and processed in R v4.3.2 using a custom script (see *Data Availability*). As some likely false positives were observed, where one dye was detected at a much later Cq value than the other detected dye, a correction was applied in R to convert the suspected false-positive heterozygote to a homozygote. Specifically, Cq differences between FAM and VIC fluorophores were calculated per well, and any cases where the absolute value of the Cq difference between the two fluorophores was greater than four, the call was determined to be a false positive for the later Cq fluorophore, which was then set to zero, making the call a homozygote for the earlier fluorophore (Figure S1 and Figure S2). Using the corrected data, genotypes were assigned based on the retained fluorophore calls: FAM only = homozygous reference; VIC only = homozygous alternate; both FAM and VIC = heterozygote; no fluorophore call = missing. Technical reproducibility was evaluated at the genotype level (Table S3), and the final call for technical replicates was based on the majority call, as inconsistent calls were re-run. Genotyping results were plotted in R (see *Data Availability*).

Statistical analysis of the Chr8 marker genotypes was carried out using IBM SPSS Statistics v28.0.1.0 (142) and R. The effect of the Chr8 marker genotype on mortality in the *Vibrio* challenge was determined using a chi-square test of homogeneity (or a Fisher’s exact test if the chi-square assumptions could not be met). To use the chi-square test, samples had to meet the following criteria: one dependent variable measured at the dichotomous level (mortality: dead vs. alive or time until death), one independent variable with three or more categorical groups (genotype: *AA, Aa, aa*), and independence of observations. Finally, the sample had to meet a minimal sample size requirement (minimal sample size for calculating expected frequencies required an expected count ≥ 5). If samples did not meet the sample size criterion, a Fisher’s exact test was performed. Baseline genotype frequencies in each family of F114-F117 prior to the *Vibrio* challenge were also assessed using this method.

### Genotyping by ddRADseq

Samples from families F114-F117, their respective parents, and OFR6-10 (VIU) were used for double-digest restriction site-associated DNA sequencing (ddRADseq) at the Institut de Biologie Intégrative et des Systems (IBIS; Quebec, Canada) using *Nsi*I/ *Msp*I cutting enzymes, as previously described (Sutherland et al., 2020). Samples were pooled to sequence 32 oysters per chip on an Ion Torrent Proton using a ThermoFisher Ion PI Chip Kit v3 chip, as per manufacturer’s instructions. Base calling was performed using Torrent Suite 5.10.1 (ThermoFisher). Multiplexed FASTQ files with sample identifying barcodes present were exported using the FileExporter plugin.

Single-end read processing and genotyping was conducted using the repository *stacks_workflow* (Normandeau, 2013/2025) and Stacks v.2.3e (Rochette et al., 2019), accompanied by custom analysis scripts (see *Data Availability*). FASTQ file quality was assessed before and after trimming using FastQC v0.11.8 with results consolidation by MultiQC (Andrews, 2010; Ewels et al., 2016). Universal adapters were eliminated and reads longer than 50 bp were retained using cutadapt v1.18 run in parallel (Martin, 2011; Tange, 2015). Quality trimming was performed on all reads to retain Phred >10 in sliding windows of size 15% of the read length, followed by read truncation to 80 bp and sample demultiplexing using the *process_radtags* module of Stacks2 (Catchen et al., 2011; Rochette et al., 2019), with two enzymes selected (i.e., *Nsi*I and *Msp*I) in parallel.

Demultiplexed and trimmed reads were aligned to a chromosome-level genome assembly for Pacific oyster (i.e., GCF_902806645.1_cgigas_uk_roslin_v1_genomic.fna) (Peñaloza et al., 2021) using bwa mem v0.7.12-r1039 (Li et al., 2009; Peñaloza et al., 2021). SAMtools v1.9 was used to remove unmapped reads and secondary or low-quality alignments (Li et al., 2009). The remaining reads following quality control were genotyped using Stacks2. The number of reads and alignments per sample, as well as alignment rates were calculated using SAMtools and custom scripts.

RAD-loci were identified with Stacks2 using the reference-based approach through the *gstacks* module with default settings. Genotype data were filtered using the *populations* module of Stacks2, requiring that a locus was genotyped in at least 70% of the individuals per population in all six populations (i.e., F114-F117, parents, and OFR6.10), and with a minimum global minor allele frequency (MAF) of 0.01. Samples with fewer than 1 M reads were identified and removed from the analysis, as previously described (Sutherland et al., 2020). Samples were genotyped again using *gstacks* as described above, and outputs were used for downstream analysis. Single SNP per RAD-tag data were exported using the *populations* module as a VCF file. The VCF file was imported into R using vcfR (Knaus & Grünwald, 2017) and converted to genind format for use with adegenet functions (Jombart & Ahmed, 2011). Functions from the *simple_pop_stats* repository were employed for analysis (see *Data availability*). The genotyping rate per sample was determined and individuals with less than 30% missing data were retained. Minor allele frequency (MAF) was re-calculated per locus, and loci with MAF < 0.01 were excluded. H_OBS_ and Hardy–Weinberg (HW) equilibrium filters were not applied due to anticipated family effects. A principal components analysis (PCA) based on genotypes was conducted using the *glPca* function of adegenet.

### GWAS for survivorship to *V. aestuarianus* challenge

A GWAS for survivorship in the *V. aestuarianus* challenge was conducted using GEMMA v0.98.5 (Zhou & Stephens, 2012) using the single-SNP per locus VCF file, as described above, converted to GEMMA input format in R (see *Data Availability*). NCBI chromosome identifiers were converted to linkage groups using the Pacific oyster assembly *Cgigas_uk_roslin_v1,* and linkage groups were converted to chromosomes following Peñaloza et al. (2021). A GWAS analysis was conducted using unimputed genotypes, as well as using genotypes that were imputed per family using the *impute_mean* function of the *missMethods* R package (Rockel, 2022). Only families F114-F117 were used for the GWAS (excluding the VIU family OFR6-10). The GEMMA analyses were conducted on families F114-F117 together by using GEMMA to calculate kinship coefficients and using either a binary phenotype (i.e., dead or alive) or a phenotype of days-to-death, where survivors of the trial were given the value of day 7 (i.e., one day after the trial concluded). A Bonferroni-corrected *p*-value of 0.05 was used as the genome-wide significance threshold. The R package *fastman* (Paria et al., 2022) was used to generate Manhattan plots from the GWAS analysis.

## Results

### Baseline genotype frequencies of Chr8 marker

Prior to the disease challenge, the Chr8 marker genotype frequencies for families F114-F117 differed from expected Mendelian proportions considering all heterozygous parents (i.e., 1:2:1 for REF/REF, REF/ALT, and ALT/ALT, respectively; Figure 1). Considering averages across the four families, heterozygotes were most frequently observed (57%), followed by the homozygous reference (25%), and the homozygous alternates were the rarest (18%). When inspecting individual families (Figure 1), a skew was observed for ALT/ALT genotypes where family F115 had significantly more than expected (35%), and F114 and F117 had significantly fewer (11% and 7%, respectively; χ^2^ test of homogeneity p<0.05). No other genotypes were significantly different among the families (Table S4).

**Figure 1.**
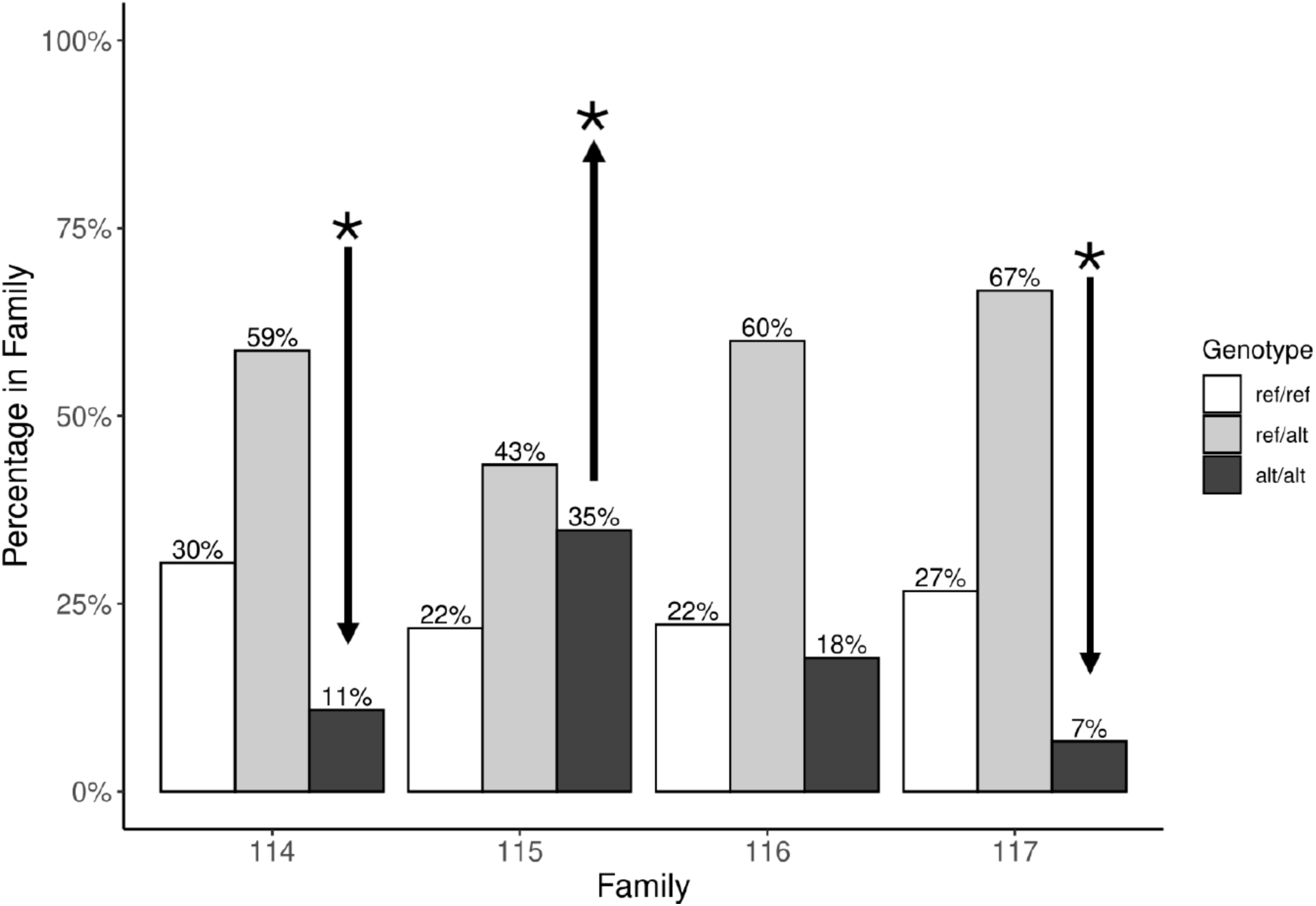
Chr8 marker genotype proportions in juveniles from families F114-F117 prior to the *Vibrio* exposure trial. Arrows indicate a genotype with a significant difference (p < 0.05) between families, where more homozygous alternate genotypes were observed in F115 relative to F114 and F117 (χ^2^=15.183, p-value = 0.019).

### *V. aestuarianus* disease challenge of all families

No mortality was observed in the first two days of the trial. Considering all the exposed, selectively bred families together (i.e., excluding the VIU breeding program OFR6-10), 37.8% of oysters died on day 3, and 67.2% died by day 6 (n = 516) (Figure S3). The remaining 252 (32.8%) survived the disease challenge. The multiple-family unexposed control group exhibited 100% survival.

The per-family cumulative mortality was on average (± s.d.) 67.2 ± 9.5% (Figure 2). Family F106 exhibited the highest mortality (85.4%), and F116 the lowest (47.9%). Rates of mortality also differed among families (p<0.001; Figure 3); F106 died more rapidly than F103, F113, F114, F115, and F116 (pairwise comparison p < 0.003; Table S5; Bonferroni correction significance threshold p = 0.003). F106 offspring were all heterozygous for the Chr8 marker, and F103 and F113 were all homozygous reference (Table S1), but when grouping families by parental genotypes at the Chr8 marker, there was no consistent cumulative mortality trend (Figure 2) and substantial inter-family variation was observed.

**Figure 2.**
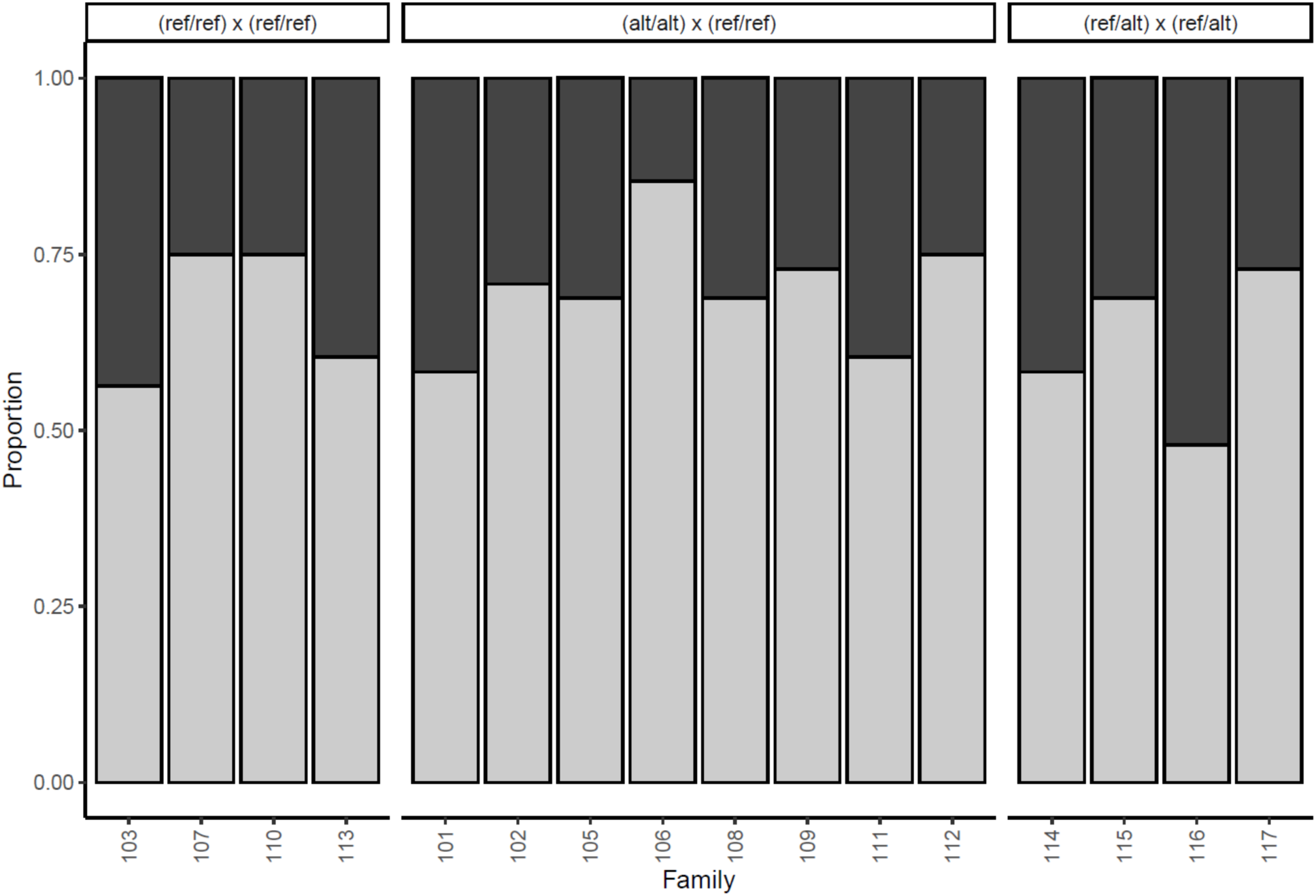
Proportions of oysters that died (light grey) or survived (dark grey) in each family in the *V. aestuarianus* disease challenge. Families are grouped by the parental genotypes at the Chr8 marker, as indicated at the top of the plot. The unexposed control group exhibited 100% survival.

**Figure 3.**
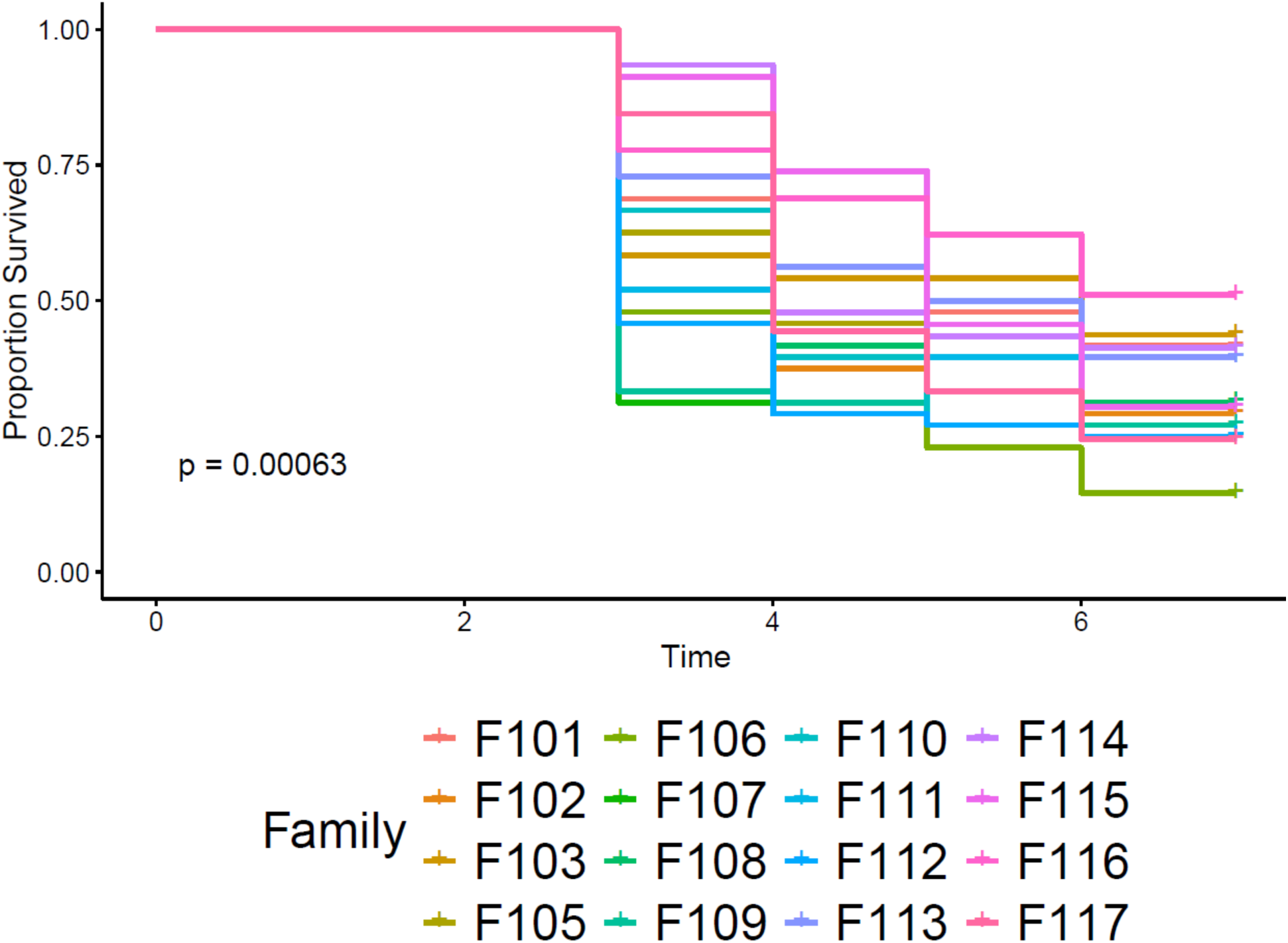
Kaplan-Meier survival curves for Pacific oyster families after exposure to *V. aestuarianus*. Survival distributions for the families showed significant differences (χ^!^=39.070, p<0.001), and pairwise differences are described in Results. Parental genotypes for the Chr8 marker per family are given in Figure 2.

### Effects of Chr8 marker allele presence and dosage on *V. aestuarianus* challenge survival

Families F114-F117 had offspring with all genotypic states of the Chr8 marker, and therefore allowed the investigation of the effects of the presence and dosage of the OsHV-1 resistance allele on mortality (days to death, Figure 4; survival, Figure 5). No significant associations were observed between offspring genotype and mortality for F114-F117 considering days to death, nor dead or alive phenotype (p>0.05). The analysis was conducted for each family individually to avoid potential confounding interfamily effects.

**Figure 4.**
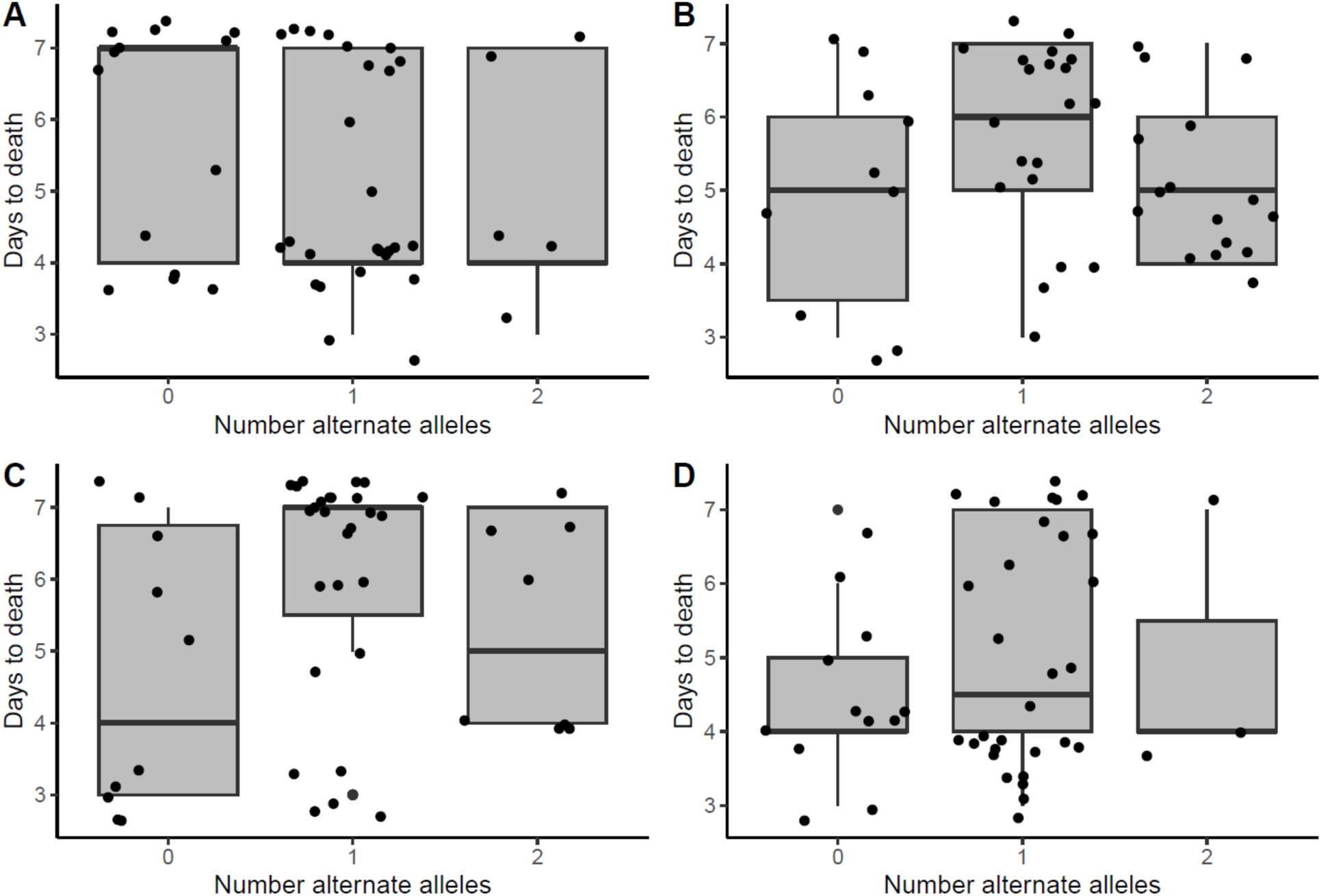
The effects of the Chr8 marker allelic dosage on mortality as quantified as days to death in the *V. aestuarianus* trial in families F114-F117 (panels A-D, respectively; i.e., those with both parents heterozygous for the Chr8 marker). No significant association was found (p>0.05) (F114 χ^!^=5.370, p>0.05; F115 χ^!^=14.172, p>0.05; F116 χ^!^=11.280, p>0.05; F117 χ^!^=4.429, p>0.05).

**Figure 5.**
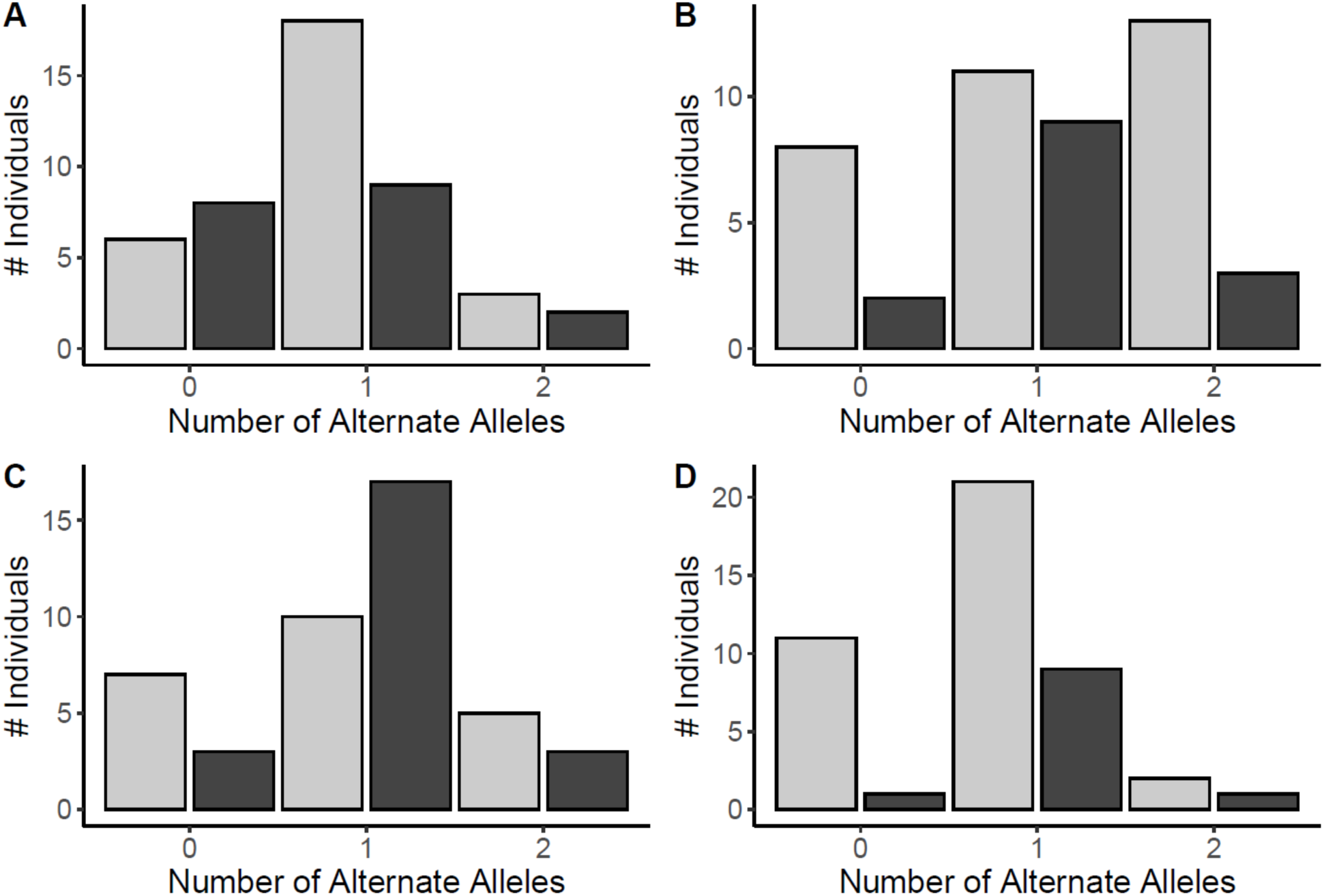
The effects of the Chr8 marker allelic dosage on mortality as quantified by the frequency of survival (light grey) or mortality (black) in the *V. aestuarianus* trial in families F114-F117 (panels A-D, respectively, i.e., those with both parents heterozygous for the Chr8 marker). No significant association was found (Chi^2^ test p > 0.05).

### ddRAD-seq genotyping of Vibrio-exposed families

Oysters from families F114-F117 and OFR6-10 (VIU family) from the *Vibrio* challenge were genotyped by ddRADseq (avg. oysters per family = 36.4). The parents of MBP families were also sequenced (Table S6). On average, 92.8 M single-end reads were produced per chip from each of the six chips sequenced (n = 523.1 M reads total). Demultiplexing resulted in an average of 91.8% reads retained. On average (± s.d.), 65.0 ± 2.2% of reads aligned against the chromosome-level reference genome (Peñaloza et al., 2021) and 340,291,536 alignments were used for genotyping with *gstacks*. In total, 277.6 M primary alignments (81.6%) were retained (10.9% of alignments were discarded owing to insufficient MAPQ and 7.5% due to excessive soft clipping). On average, 1.79 M alignments were obtained per sample (range: 0.265-4.681 M records per sample). The removal of samples with fewer than 1 M reads resulted in the removal of 18 of the 190 samples, retaining 172 samples (Table S6). Notably, 15 of these 18 removed samples were oysters that had died during the trial, suggesting that these samples may have had substantial DNA degradation occurring between the mortality event and sampling, which has also seen in other disease challenge trials in Pacific oysters (Sutherland et al., 2025).

After low-yield samples were removed, genotyping used 330.6 M alignments (269.7 M primary alignments) to build 393,153 RAD loci with an average (± s.d.) effective coverage of 56.9 ± 22.6x. Low genotyping rate loci were removed (see Methods), as were those with MAF < 0.01, resulting in the retention of 10,202 RAD loci comprising 889,291 genomic sites, of which 25,801 contained variants (percent polymorphic = 2.9%). The exported VCF file with a single SNP per RAD locus retained resulted in 8,252 variants for downstream analysis. Low genotyping rate samples (missing ≥ 30% of loci) were removed, retaining 168 individuals (Figure S4; Table S6), including 32, 36, 35, 33, and 25 individuals from F114, F115, F116, F117, and OFR6-10, respectively, as well as seven of the eight parents of families F114-F117. Reapplying the MAF filter resulted in the retention of 8,193 SNPs (4,218 with MAF > 0.1). Clustering samples based on genotypes by PCA showed close clustering of samples within the four MBP families, with the parents of these families interspersed among the offspring and the VIU family OFR6-10 grouping together but apart from the MBP families on PC1 (Figure S5). One individual labeled as F116 clustered within the F115 family, suggesting a sample labelling or handling error (Figure S5).

### GWAS for *V. aestuarianus* survival

Selectively bred oysters (families F114-F117) were used in GWAS analyses considering survival to the *V. aestuarianus* challenge as a binary response (dead or alive) or days-to-death. This included 49 oysters that survived and 87 that died (total n = 136). Considering only loci on chromosomes (n = 7,870 SNPs), without imputation 3,656 SNPs were analyzed by GEMMA with a MAF > 0.05 filter applied. The percent variation explained (PVE) ± standard error (S.E.) was 19.2 ± 14.7% for survival as a binary trait and 6.3 ± 9.8% for days post-exposure survival. There were no significantly associated loci to survivorship to the *Vibrio* challenge considering survival as dead or alive, or as days-to-death (Figure 6). Similarly, no significant associations were identified using the day-to-death phenotype (Figure S6A). Using mean imputed genotypes, 5,478 of the 7,870 chromosomal SNPs were analyzed by GEMMA (MAF > 0.05), resulting in similar PVE values (PVE ± S.E. for dead/alive variable = 18.1 ± 14.8% and for days-to-death = 5.7 ± 9.6%). The GWAS with imputed genotypes also did not identify any loci significantly associated with the phenotype survival or days-to-death (Figure S6B and Figure S6C).

**Figure 6.**
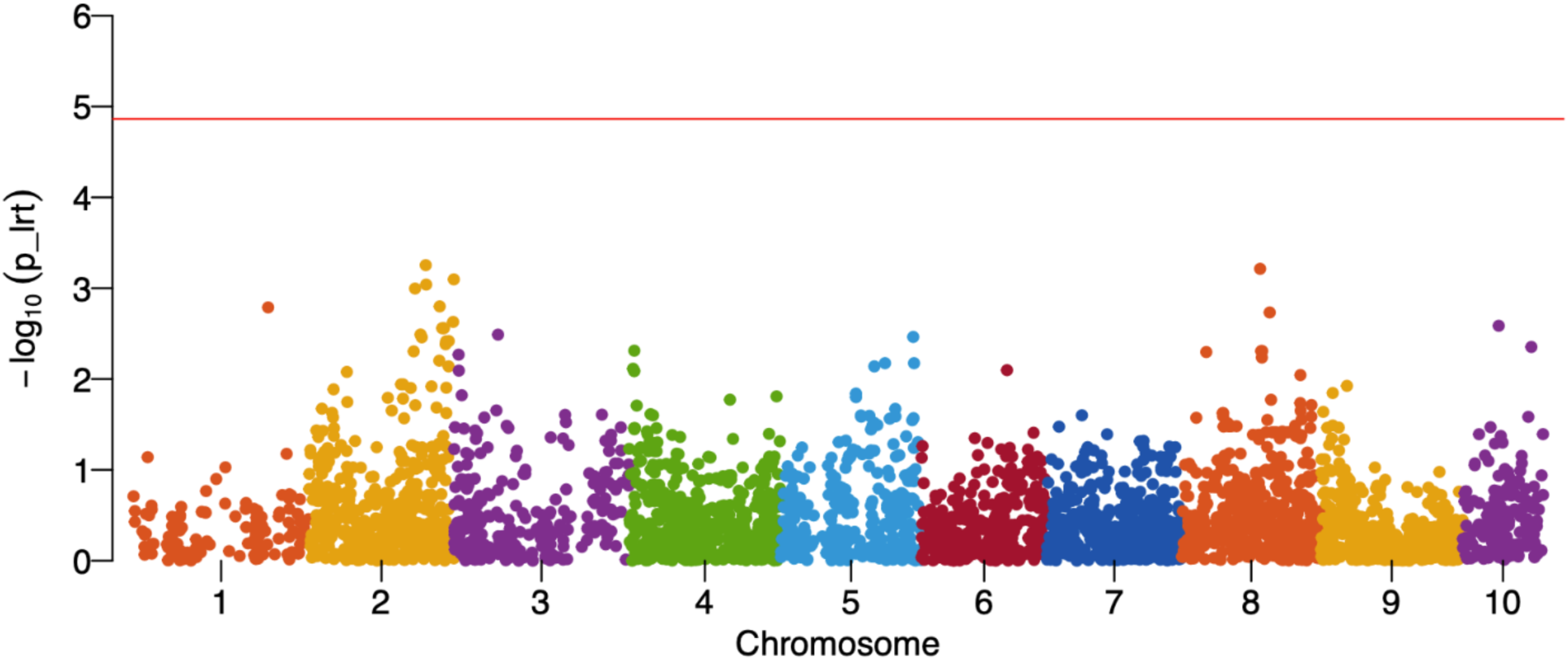
Manhattan plot showing associations of each SNP from the unimputed RAD-seq dataset for survivorship (mortality or survival) in the *V. aestuarianus* trial for families F114-F117. The red horizontal line indicates the genome-wide Bonferroni-corrected p-value significance threshold of 0.05. No significant associations were observed.

## Discussion

Host genetic and epigenetic factors are known to influence disease dynamics (Krickeberg et al., 2012; Magwire et al., 2012). In this study, we assessed whether the presence of a marker associated with increased survival and resistance to a virus (OsHV-1) affects susceptibility to a bacterial pathogen (*V. aestuarianus*). Our results indicate that the presence and allelic dosage of the OsHV-1 resistance allele of the Chr8 marker (Divilov et al., 2023) in selectively bred families of Pacific oysters does not affect the susceptibility of the juvenile oysters (spat) to *V. aestuarianus* infection. This contrasts with recent findings that the marker in many of these same families is associated with increased survivorship to *V. coralliilyticus* infection (Lunda et al., 2026). The cause of the different responses to the two bacteria species is not known, since both *V. coralliilyticus* and *V. aestuarianus* would potentially be present in the environment in which the field trials for survivorship occurred, and therefore the selective advantage of the surviving oysters could have captured increased resistance to the bacteria in addition to the endemic OsHV-1, as discussed for *V. coralliilyticus* by Lunda et al. (2026).

Juvenile Pacific oysters are susceptible to *V. aestuarianus* in British Columbia (BC), Canada (Cowan et al., 2024). The present results and the results of Lunda et al. (2026) provide important insights for selective breeding for disease resistance in Pacific oysters. The present results suggest a lack of pleiotropy in the Chr8 marker for *V. aestuarianus* resistance and an absence of genetic correlation between survivorship to OsHV-1 and *V. aestuarianus*, at least in regard to the survivorship mechanism associated with the Chr8 marker. The results of Lunda et al. (2026) indicate that there should be no negative consequence of using the Chr8 marker for marker-assisted selection on the resistance of the selected oysters for *V. coralliilyticus*, and in contrast should improve their resilience to that bacterial pathogen. Given the independence observed between the Chr8 protective allele and *V. aestuarianus* resistance, identifying loci involved in resistance to *V. aestuarianus* may enable marker-assisted selection of oysters that are resistant to both OsHV-1 (and *V. coralliilyticus*; Lunda et al., 2026) and *V. aestuarianus*. However, in the present study we were not able to identify any major effect loci for resistance to *V. aestuarianus* and rather identify that resistance may be controlled by polygenic architecture.

### Effects of the OsHV-1 resistance allele on survival to *Vibrio aestuarianus* infection

The presence of the Chr8 marker allele associated with increased resistance to OsHV-1 had no effect on the survival to *V. aestuarianus* infection. This finding is supported by another study that reported the absence of genetic correlations between OsHV-1 and *V. aestuarianus* resistance (Azéma et al., 2017). Azéma et al. (2017) analyzed the genetic parameters for resistance to OsHV-1 and *V. aestuarianus* at three different life stages: spat (3 to 6 months old); juvenile (11 to 15 months old); and adult (25 to 26 months old). They reported no observable genetic correlation between survival to viral or bacterial infection. However, the life stage of the oyster significantly affected susceptibility to both pathogens; susceptibility to OsHV-1 decreased with age and growth, whereas susceptibility to *V. aestuarianus* increased with age and growth. The different effects of age on susceptibility to OsHV-1 and *Vibrio* may reflect the different immune responses of oyster hosts. Further evidence of the independence of the resistance mechanisms was provided by Green et al. (2016), who found different transcriptome responses to OsHV-1 and *V. aestuarianus* infections. Thus, the immune genes involved in the defence against OsHV-1 and *Vibrio aestuarianus* infections appear to be distinct.

The absence of a correlation between the Chr8 marker and *V. aestuarianus* susceptibility suggests that the mechanisms of resistance to these pathogens are independent. The Chr8 marker allele associated with increased survivorship is thought to upregulate *interferon regulatory factor 2* (*irf2*), an antiviral gene that can provide oysters with constitutively upregulated antiviral immunity (Divilov et al., 2023). With constitutively overexpressed *irf2*, overexpression of antiviral genes could be expected even in the absence of the virus. Therefore, the lack of increased susceptibility to *V. aestuarianus* is notable, as it suggests that the upregulated antiviral immunity does not impact (negatively or positively) the oyster’s ability to survive the bacterial infection. However, the finding of increased resistance to *V. coralliilyticus* (Lunda et al., 2026) suggests that this is not a universal conclusion, and that the mechanism of resistance does provide resistance to a bacteria pathogen (i.e., *V. coralliilyticus*). The genetic mechanisms underlying *Vibrio* resistance are not well known. However, a recent study using 52 Pacific oyster families found 18 SNPs in three QTL associated with resistance to *V. alginolyticus* in *C. gigas,* and the authors suggested that *Vibrio* resistance is polygenic (Yang et al., 2022). Most of the associated genes are related to pathogen recognition and immune defence responses (Yang et al., 2022). Collectively, genes that induce the expression of immune-related genes are potential targets for selective breeding to increase *Vibrio* resilience.

Selective breeding for one trait can lead to fitness trade-offs by negatively affecting a second trait and consequently impacting overall fitness (Divilov et al., 2024; Dwivedi et al., 2021). Our understanding of the compromises in selection for disease resistance is limited, particularly regarding the effects on other traits or resistance to other pathogens. Trade-offs can arise when beneficial traits are linked to detrimental ones, thereby affecting fitness outcomes (Gallardo-Hidalgo et al., 2021). For example, although selection for increased growth rate has also improved larval resilience to ocean acidification, fast-growing oysters are more susceptible to marine heatwaves and *V. aestuarianus* infections later in life; mortality has been found to increase by 76% per gram of body weight gained (Khtikian, 2021; Nordio et al., 2021). However, selection for OsHV-1 resistance in the MBP breeding population has not been observed to impact growth (Divilov et al., 2024). Selection for disease resistance is complex, as resistance is expected to be influenced by multifactorial biological functions, including reproduction and immunity (Bai & Plastow, 2022; Wang et al., 2021).

We expected that a trade-off could be observed in the present study because up-regulation of the immune system can increase the energy requirements of the host (Lochmiller & Deerenberg, 2000; Robinson & Green, 2020). This can reduce the overall reproductive success, growth rates, and survival to events such as pathogen outbreaks (Lochmiller & Deerenberg, 2000; Robinson & Green, 2020). Additionally, in some cases, over-reactive immune responses to pathogens can be more damaging than a weak immune response (Green et al., 2016), as discussed in relation to immune tolerance (Råberg et al., 2007). The expression of extremely potent antibacterial or antiviral effectors can result in a toxic cellular environment that can be detrimental to the organism (Green et al., 2016), and strong immune effectors and responses can inflict damage on host tissues. In the case of infectious diseases, a fitness trade-off might occur if the heightened immune response leads to greater collateral damage (Råberg et al., 2007). Furthermore, other environmental stress factors may impact immune responses due to energetic constraints (Sutherland et al., 2014). Here, we did not measure other factors that may represent energetic trade-offs for increased viral resistance (e.g., growth rate and response to other stressors), but these factors could be integrated into future studies to evaluate the impact of the Chr8 marker on *Vibrio* resistance. Notably, before the *V. aestuarianus* exposure occurred, there were baseline differences in the genotype frequencies of the Chr8 marker, specifically in the proportion of the homozygous alternate genotype. Therefore, other unknown factors may cause distortions in the frequency of this marker, which could affect future breeding practices around the Chr8 marker.

Although we did not observe a trade-off, selection for disease resistance may also indirectly increase resistance to other stresses such as marine heat waves (MHW). MHWs increase water temperature, resulting in high mortality rates (Cowan et al., 2024). Vibriosis and *V. aestuarianus* proliferation are further linked to seawater temperature, with increased temperature increasing host susceptibility by weakening the immune system and increasing the transmission and proliferation of pathogenic *Vibrio* species (Cowan et al., 2024; Wendling et al., 2014). MHWs are a consequence of climate change, which is driven by increased anthropogenic emissions of CO_2_. The global average MHW frequency and duration increased by 34% and 17%, respectively, resulting in a 54% increase in annual marine heatwave days globally (Oliver et al., 2018). Bolstering resilience to MHWs may be achieved by producing oysters with increased resistance to viral and bacterial pathogens. This can mitigate the detrimental effects of these temperature spikes on oyster populations. This dual-pathogen resistance strategy could serve as a multifaceted approach to bolster oyster health and ensure the sustainability of oyster aquaculture in the presence of various environmental stressors.

### Family effects and GWAS for *V. aestuarianus* resistance

In this study, we also analyzed family differences in survival following *V. aestuarianus* infection, both in cumulative mortality and mortality rates (days to death). Although different families have known OsHV-1 genotypes, they also would have other genetic differences that could affect resistance to *V. aestuarianus*. As the Chr8 marker did not explain the variance observed across families in *Vibrio* susceptibility, other factors may be involved.

In the GWAS analysis for survivorship to *V. aestuarianus*, we did not identify any loci significantly associated with mortality or mortality rate in response to infection. The applied sample size may have been too small to detect a more subtle genetic association, such as that which would occur with a polygenic trait. Resistance to *Vibrio* infections has previously been reported to exhibit a polygenic architecture (Yang et al., 2022), and therefore may require a significantly larger sample size to identify significant associations. It is also possible that inspecting other populations of Pacific oysters could identify major effect loci for *Vibrio* survivorship that are not present in the MBP families investigated here. Furthermore, differences in resistance across families could involve other determining factors, such as epigenetic differences impacting immune responses to the bacterial infection. For example, a recent GWAS for survival to Pacific oyster mortality syndrome indicated that epigenetic factors play a larger role than genetic factors for the syndrome (Gawra et al., 2023). Future studies may benefit from including a larger number of individuals to account for the potential polygenic nature of *Vibrio* resistance mechanisms, but also may need to consider that *Vibrio* resistance as a trait could be best addressed by other selective breeding approaches, including genomic selection (Boudry et al., 2021; Meuwissen et al., 2016), rather than marker-assisted selection, in part because of the challenge in identifying major effect loci.

## Conclusion

The genotype and allelic dosage of the Chr8 marker allele associated with survivorship to OsHV-1 did not affect susceptibility to *V. aestuarianus* infection in selectively bred Pacific oyster spat evaluated here. This suggests a lack of pleiotropic effect of the Chr8 marker on *V. aestuarianus* resistance, as well as a lack of genetic correlation between these traits. Given the recent findings that the OsHV-1 resistance allele also provides protection to *V. coralliilyticus*, this may suggest different immune pathways for defense against these two *Vibrio* species. The GWAS analysis did not find any loci significantly associated with *Vibrio* survivorship, suggesting a polygenic architecture in these families for this trait. Nonetheless, significant family differences in *Vibrio* susceptibility were observed here, even within the MBP breeding program families that have been selectively bred for at least six generations (de Melo et al., 2016) and therefore are expected to be more genetically homogenous than outbred oysters. These observations emphasize the potential for further investigation into selective breeding for resistance to *V. aestuarianus*, which may benefit from genomic selection rather than marker-assisted selection, unless major effect loci can be identified.

## Supplementary Materials

Supplementary tables and figures are provided in Supplementary Results.

**Additional File S1.** Read counts and alignments for all samples retained in the analysis.

## Data Availability

Raw sequencing data have been uploaded to SRA under BioProject PRJNA1145972 with BioSamples SAMN43080749 - SAMN43080938.

RADseq genotypes and supporting files: 10.6084/m9.figshare.26524321

rhAmp assay raw data plate files: 10.6084/m9.figshare.26515438

Sample and family IDs for mapping families: 10.6084/m9.figshare.28599923

This analysis was supported by the following repositories:

Manuscript code and README: https://github.com/bensutherland/ms_cgig_chr8_vibrio

simple_pop_stats repository: https://github.com/bensutherland/simple_pop_stats

## Funding

Funding for this study was provided by the Pacific States Marine Fisheries Commission (PSMFC) project “Preparing for Future Challenges – Threats from Ocean Acidification, *Vibrio coralliilyticus* and OsHV-1 μvar to West Coast Oyster Farmers” (Award No. NA18NMF4720007).

## Conflicts of Interest

BJGS is affiliated with Sutherland Bioinformatics. The other authors declare no conflicts of interest.

## Acknowledgements

Thanks to the members of the Centre for Shellfish Research of Vancouver Island University for laboratory assistance, to Antonia Barela, Gary Fleener, and William Schoeneck for assistance in the hatchery for the production of juvenile oysters, and to the Deep Bay Marine Field Station staff for storing and acclimating the juvenile oysters. Thanks to Brian Boyle at IBIS for ddRADseq library preparations and sequencing. Thanks to three anonymous reviewers for comments on an earlier version of this manuscript.

## Author Contributions

BS, LS, TG, and CL conceived of the study. LS, SL, TG conducted laboratory experiments. BS, LS, and KD contributed to software development and analysis. BS curated the data. LS and BS drafted the original manuscript. TG and BS provided laboratory and staff supervision. TG and CL acquired funding for the work and administered the project. All authors (LS, BS, SL, AL, KD, CL, TG) supported result interpretation and contributed to manuscript review and editing.

## Notes

https://github.com/bensutherland/ms_cgig_chr8_vibrio

https://doi.org/10.6084/m9.figshare.26515438.v1

https://doi.org/10.6084/m9.figshare.26524321

### Summary of Updates

Revisions added following the publication of Lunda et al. (2026) to ensure we mention the different results observed in the two works.

