## Supplemental Information for "The presence of a marker associated with Pacific oyster resistance to OsHV-1 does not affect susceptibility to *Vibrio aestuarianus*"

### Table of Contents


| | |
|---|---|
| **Figure S1. rhAmp assay false-positive heterozygote correction in families F114-F117** | **Page 2** |
| **Figure S2. rhAmp assay false-positive heterozygote correction in families F101-F103 and F105-F112.** | **Page 3** |
| **Figure S3. Daily mortality of *C. gigas* during the disease challenge** | **Page 4** |
| **Figure S4. ddRADseq genotyping rate per individual** | **Page 4** |
| **Figure S5. PCA of ddRADseq SNP data** | **Page 5** |
| **Figure S6. Manhattan plots for different iterations of the GWAS analysis, including days-to-death and with or without mean imputation of genotypes.** | **Page 6** |
| **Table S1. Chr8 family crosses and parental genotypes** | **Page 8** |
| **Table S2. rhAmp assay PCR conditions** | **Page 8** |
| **Table S3. rhAmp assay interplate and intraplate variation** | **Page 9** |
| **Table S4. Summary of genotype proportions per family.** | **Page 9** |
| **Table S5. Pairwise log-rank comparisons for differences in mortality for F106** | **Page 9** |
| **Table S6. Summary of samples sequenced by ddRADseq** | **Page 10** |
| **Additional results. rhAmp assay reproducibility and plate variation and DNA quality of all families** | **Page 11** |

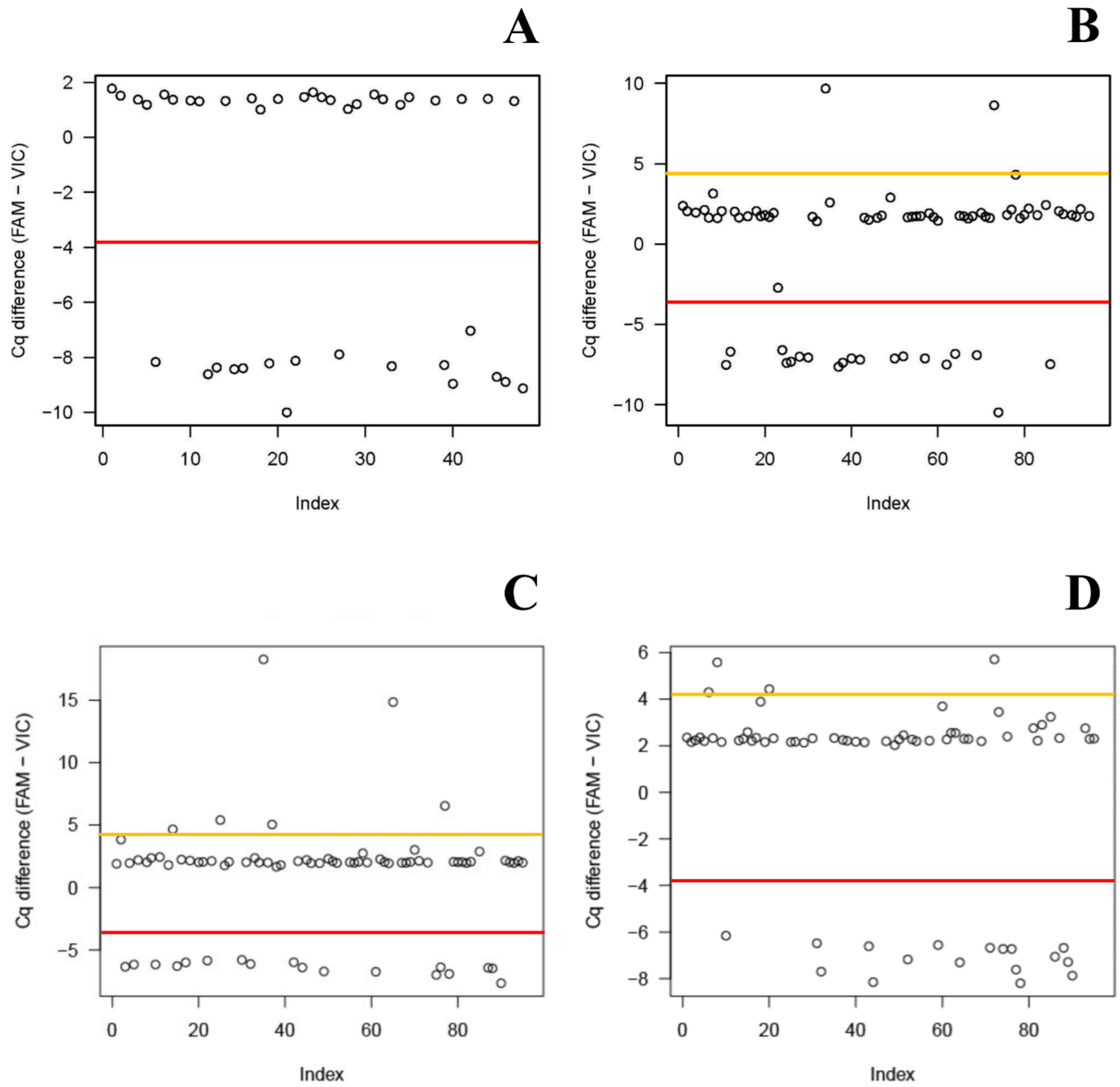


**Figure S1.** Method of correcting false positive heterozygotes in rhAmp assay for individuals in families F114-F117. The difference between the Cq values for VIC and FAM fluorophores was determined and plotted per plate (panels A-D show plates 1-4, respectively). Differences between the fluorophores that were greater than or equal to +4 were deemed to be false positive FAM and subsequently corrected and set to 0. Differences less than or equal to -4 were deemed to be false positive VIC and subsequently corrected and set to 0. Samples above the orange line indicate false-positive FAM detections, while samples below the red line indicate false-positive VIC detections. Samples that are above the red line and below the yellow line indicate true-positive heterozygotes.

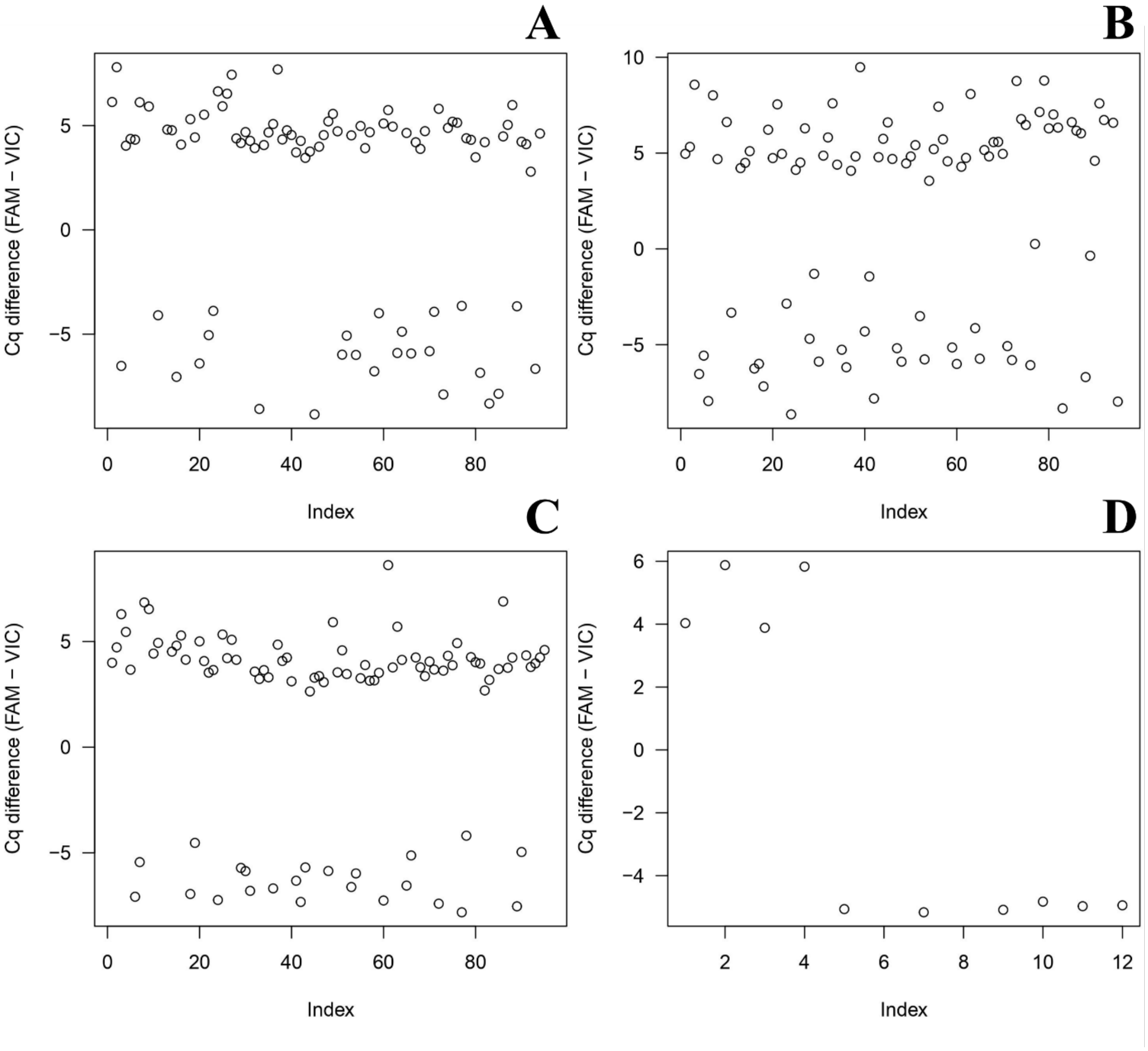


**Figure S2.** Method of correcting false positive heterozygotes in rhAmp assay for individuals in families F101-F103 and F105-F112. The difference between the Cq values for VIC and FAM was determined and plotted per plate (panels A-D show plates 6-9, respectively). Due to the lack of normalization, it was difficult to set cutoffs to detect false positive heterozygotes with the extracted DNA.

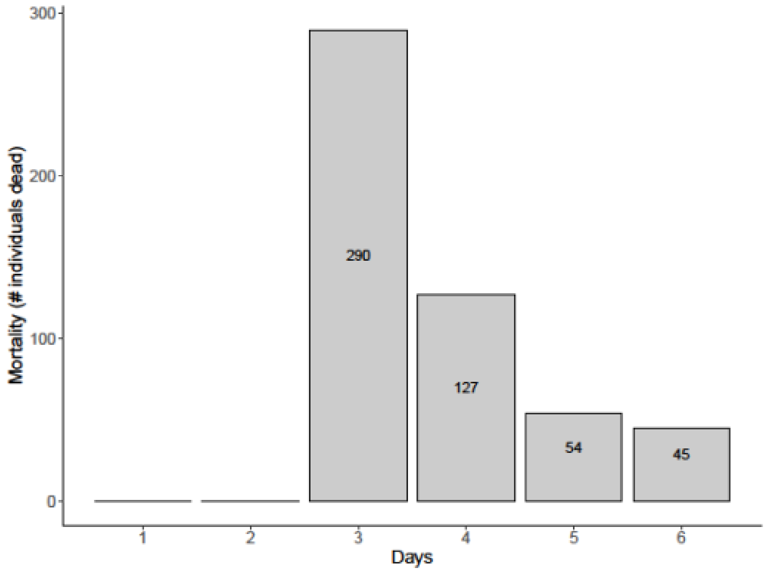


**Figure S3.** Daily mortality of *C. gigas* across all families (F101-F103, F105-F117) after exposure to *V. aestuarianus* during the disease challenge experiment. No mortality was observed on days 1 and 2.

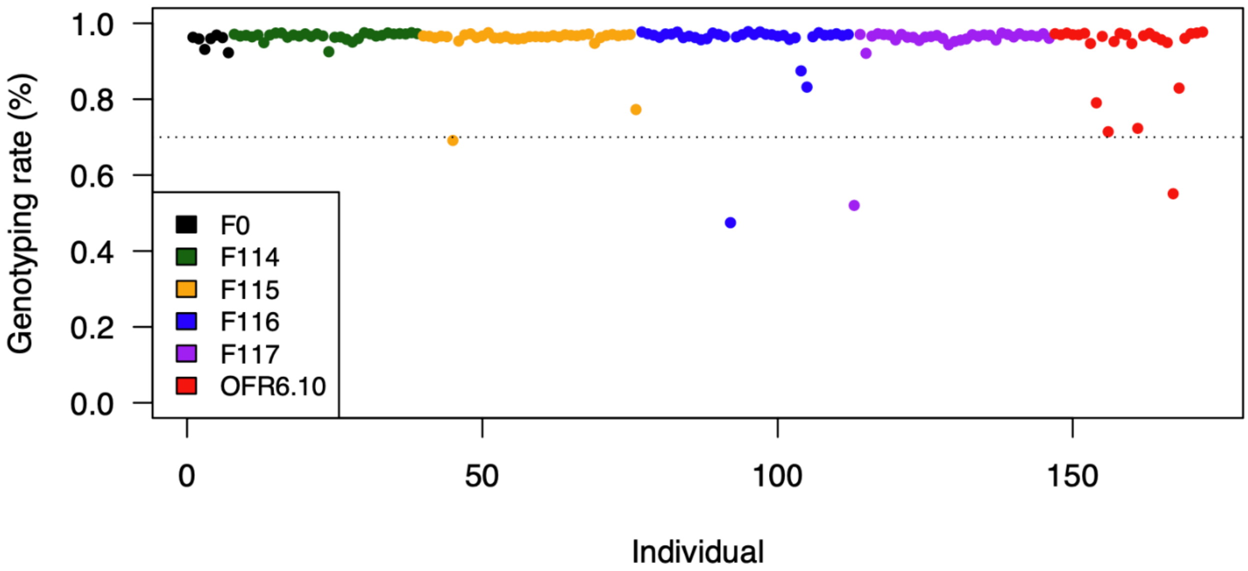


**Figure S4.** Genotyping rate per individual coloured by family. F 114 through F117 are the mapping families from the selectively bred OsHV-1 resistant population, and OSU.F0 are the parents of these families. OFR6.10 is the VIU breeding program family. A filter was subsequently applied to remove samples with genotyping rate < 70%, as indicated by the horizontal dotted line.

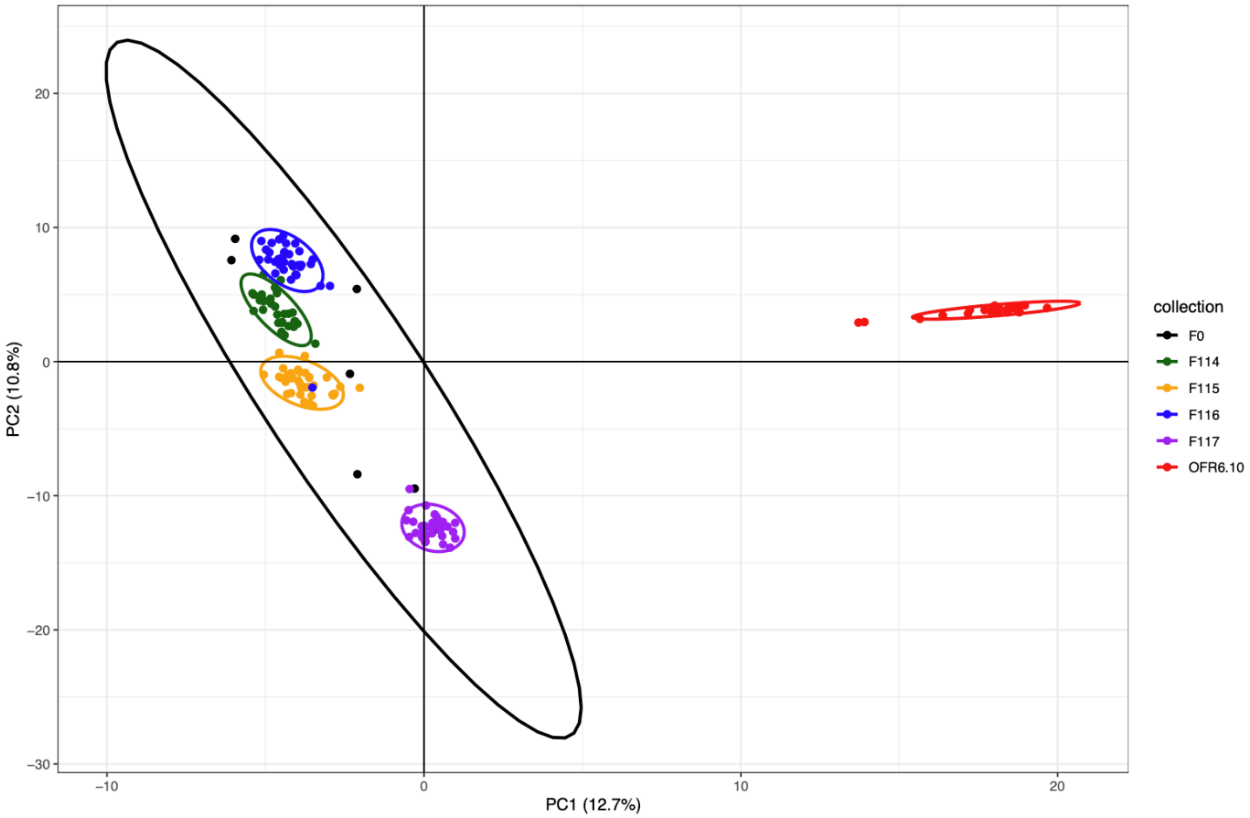


**Figure S5:** Principal component analysis (PCA) of SNP data generated from ddRADseq showing PC1 and PC2. Data points are labelled by family/group. Families/groups are organized based on colour PC1 explained 12.7% of the data, while PC2 explained 10.8%. The VIU family (OFR6.10) shows the most differentiation from the OSU families (F114, F115, F116, F117,) and the OSU parents (OSU.F0).

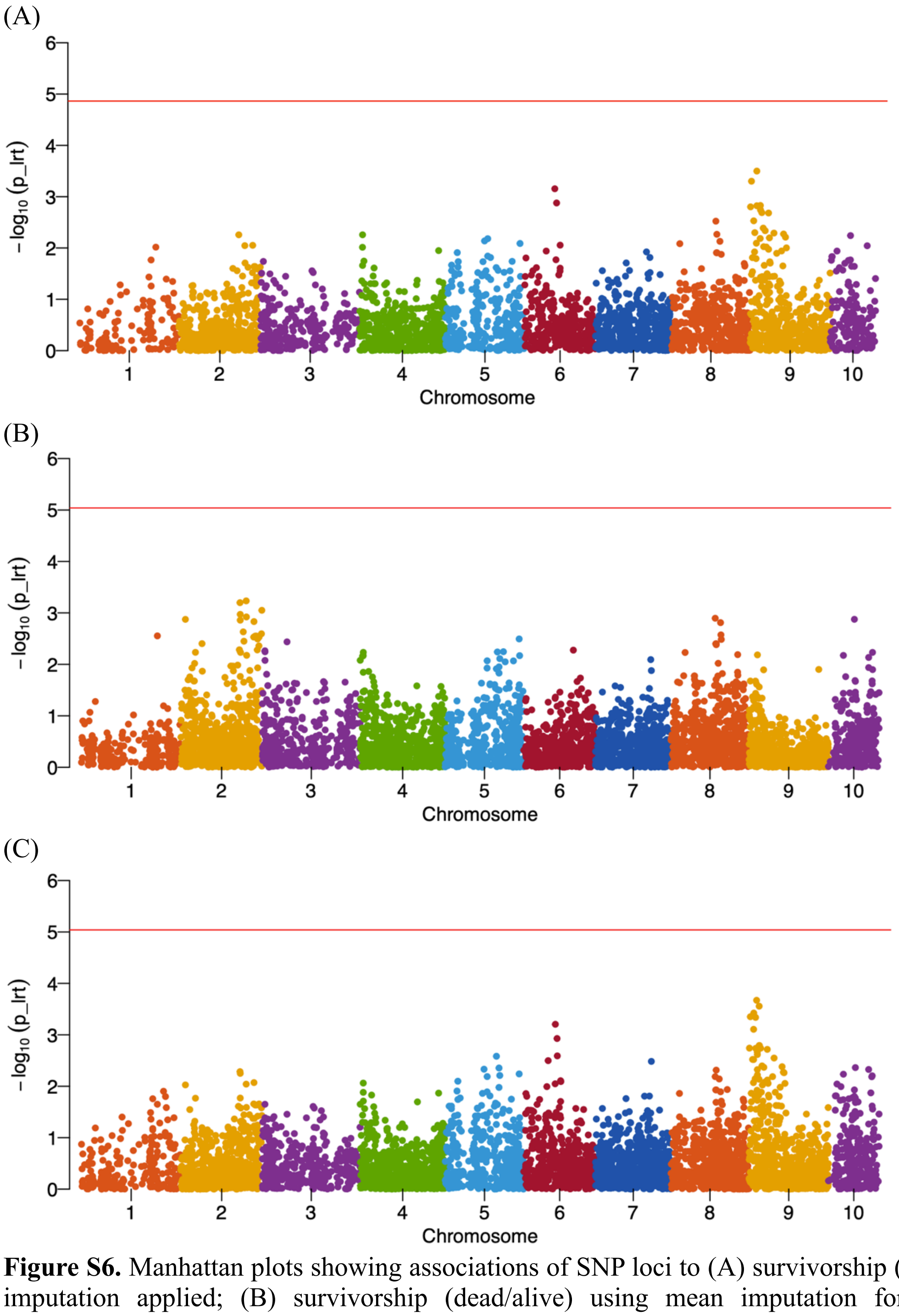


**Figure S6.** Manhattan plots showing associations of SNP loci to (A) survivorship (days-to-death) without imputation applied; (B) survivorship (dead/alive) using mean imputation for genotypes; and (C) survivorship (days-to-death) using mean imputation for genotypes. All MBP families (i.e., F114-F117) were analyzed together, using a kinship matrix as calculated by GEMMA. The red horizontal line represents the genome-wide Bonferroni-corrected p-value significance threshold of 0.05. No significant associations were observed.

**Table S1.** Summary of Chr8 crosses and parental genotypes at the OsHV-1 resistance marker. The alternate allele (alt) is the allele with protective phenotype against OsHV-1, and the reference allele (ref) is the more frequent allele in nature.

| Family | Parent 1 genotype | Parent 2 genotype |
|---|---|---|
| 101 | (alt/alt) | (ref/ref) |
| 102 | (alt/alt) | (ref/ref) |
| 103 | (ref/ref) | (ref/ref) |
| 104 | (ref/ref) | (ref/ref) |
| 105 | (alt/alt) | (ref/ref) |
| 106 | (alt/alt) | (ref/ref) |
| 107 | (ref/ref) | (ref/ref) |
| 108 | (alt/alt) | (ref/ref) |
| 109 | (alt/alt) | (ref/ref) |
| 110 | (ref/ref) | (ref/ref) |
| 111 | (alt/alt) | (ref/ref) |
| 112 | (alt/alt) | (ref/ref) |
| 113 | (ref/ref) | (ref/ref) |
| 114 | (ref/alt) | (ref/alt) |
| 115 | (ref/alt) | (ref/alt) |
| 116 | (ref/alt) | (ref/alt) |
| 117 | (ref/alt) | (ref/alt) |

**Table S2.** Summary of rhAmp assay PCR conditions.

| Step | Temperature (ºC) | Duration | Cycles |
|---|---|---|---|
| Enzyme activation | 95 | 10min | 1 |
| Denaturation | 95 | 10s | |
| Annealing | 60 | 30s | 40 |
| Extension | 68 | 20s | |
| Heat inactivation | 99.9 | 15min | 1 |

**Table S3.** Summary of inter-plate and intra-plate variation represented as standard deviation (±Cq)

| | Standard Deviation (± Cq) | |
|---|---|---|
| | FAM | VIC |
| Intraplate variation (between wells) | 0.20 Cq | 0.24 Cq |
| Interplate variation (between plates | 0.69 Cq | 0.46 Cq |

**Table S4.** Summary of genotypes per family. Asymptotic significance from chi-square test: p<0.05. Each subscript letter denotes a subset of Family categories whose column proportions do not differ significantly from each other at the .05 level.

| Genotype | | Family | | | | |
|---|---|---|---|---|---|---|
| | | **F114** | **F115** | **F116** | **F117** | **Total** |
| Homo.ref (ref/ref) | Count | $14_a$ | $10_a$ | $10_a$ | $12_a$ | 46 |
| | % within Family | 30.4% | 21.7% | 22.2% | 26.7% | 25.3% |
| Het (ref/alt) | Count | $27_a$ | $20_a$ | $27_a$ | $30_a$ | 104 |
| | % within Family | 58.7% | 43.5% | 60.0% | 66.7% | 57.1% |
| Homo.alt (alt/alt) | Count | $5_a$ | $16_b$ | $8_{a, b}$ | $3_a$ | 32 |
| | % within Family | 10.9% | 34.8% | 17.8% | 6.7% | 17.6% |

**Table S5:** Pairwise log-rank comparisons for differences in mortality for F106. Significance was determined after Bonferroni correction (p< 0.003)

| Family compared to F106. | $\chi^2$ | p-value |
|---|---|---|
| F103 | 9.340 | 0.002 |
| F113 | 9.292 | 0.002 |
| F114 | 14.017 | p<0.001 |
| F115 | 11.149 | p<0.001 |
| F116 | 16.828 | p<0.001 |

**Table S6.** The number of samples sequenced by ddRADseq, then passing a filter of low reads (retain when more than 1 M demultiplexed reads per sample), and a filter on genotyping rate per sample of at least 70% of the RAD-loci.

| Group | Samples sequenced | Avg. reads (M) | Samples >1 M reads | Samples GR ≥ 70% | Trial survivors | Trial mortalities |
|---|---|---|---|---|---|---|
| MBP parents | 8 | 2.5 | 7 | 7 | - | - |
| F114 | 36 | 2.8 | 32 | 32 | 14 | 18 |
| F115 | 38 | 2.9 | 37 | 36 | 10 | 26 |
| F116 | 40 | 2.8 | 36 | 35 | 18 | 17 |
| F117 | 37 | 2.5 | 34 | 33 | 7 | 26 |
| OFR6.10 | 31 | 2.7 | 26 | 25 | 14 | 11 |
| ***Total*** | **190** | | **172** | **168** | **63** | **98** |

**Technical analysis: rhAmp assay reproducibility and plate variation.**

Reproducibility and variation were assessed at the genotype level, the plate level, and between individual PCR wells. When comparing duplicates, 175 samples (95%) were completely concordant in genotype calls. Nine samples were discordant. For the discordant samples, the genotype was assigned correctly 63% of the time. Reproducibility and variation among plates and wells were determined using standard deviation (The standard deviation calculated for intraplate variation for FAM and VIC was 0.20 quantification cycles (Cq) and 0.24 Cq, respectively, between wells on each plate). The standard deviation calculated for interplate variation for FAM and VIC was 0.69 Cq and 0.49 Cq, respectively, showing a relatively low deviation between both individual wells and individual plates and indicating adequate reproducibility throughout the study

**Technical analysis: DNA quality of *V. aestuarianus* exposed oysters (all families)**

Quality control and quantification of column extractions of genomic DNA was performed using absorbance measurements, Qubit fluorimetry, and gel electrophoresis. The average column extraction concentration measured with absorbance was 130.0 ± 77 ng/uL (SD). The average column extraction concentration from Qubit fluorimetry was 69.3 ± 59 ng/uL (SD). The average A260/A280 ratio measured with absorbance was 1.88.0 ± 0.19 (SD) while the average A260/A230 was 1.76 ± 0.55 (SD). The percent difference between the average concentration measured by absorbance versus the average concentration measured by Qubit fluorimetry was -87.7%. Gel electrophoresis with a 1% agarose gel was used to assess DNA for quality. Quality was determined based on observable smearing which indicates degraded DNA. Furthermore, quality was assessed based on how much DNA pooled at a position matching the lowest band of the 1 kbp ladder, which indicates severity of degradation. Concentration ratings were based on the measured absorbance values (low = ≤75ng/uL, medium =>75ng/uL≤150ng/uL, high = >150ng/uL). Additionally, death day and extract day were analyzed to see their effect on quality with regards to DNA degradation; however, no significant association was found between these variables and DNA quality.
